# Streptolysin production and activity is central to *in vivo* pathotype and disease outcome in GAS infections

**DOI:** 10.1101/686303

**Authors:** Jenny Clarke, Murielle Baltazar, Mansoor Alsahag, Stavros Panagiotou, Marion Pouget, William A Paxton, Georgios Pollakis, Dean Everett, Neil French, Aras Kadioglu

## Abstract

Streptococcus pyogenes (GAS) is among the most diverse of all human pathogens, responsible for a range of clinical manifestations, from mild superficial infections such as pharyngitis to serious invasive infections such as necrotising fasciitis and sepsis. The drivers of these different disease phenotypes are not known. The GAS cholesterol-dependent cytolysin, streptolysin O (SLO), has well established cell and tissue destructive activity. We investigated the role of SLO in determining disease outcome *in vivo*, by using two different clinical lineages; the recently emerged hypervirulent outbreak *emm* type 32.2 strains, which result in sepsis, and the *emm* type 1.0 strains which cause septic arthritis. Using clinically relevant *in vivo* mouse models of sepsis and a novel septic arthritis model, we demonstrated that the amount and activity of SLO is vital in determining the pathotype of infection. The *emm*32.2 strain produced large quantities of highly haemolytic SLO that resulted in rapid development of sepsis. By contrast, the lower levels and haemolytic activity of *emm*1.0 SLO led to translocation of bacteria to joints. Importantly, sepsis associated strains that were attenuated by deletion or inhibition of SLO also translocated to the joint, confirming the key role of SLO in determining infection niche. Our findings demonstrate that SLO is key to *in vivo* pathotype and disease outcome. Careful consideration should be given to novel therapy or vaccination strategies that target SLO. Whilst neutralising SLO activity may reduce severe invasive disease, it has the potential to promote chronic inflammatory conditions such as septic arthritis.

## Introduction

Group A *Streptococcus* (GAS), also called *Streptococcus pyogenes*, is a commensal of the human upper respiratory tract and also an important human pathogen, accounting for over 750 million infections every year ^1,2^. GAS is able to produce a variety of pyogenic infections that range in severity and prevalence ^3-5^. Diseases include pharyngitis, impetigo, cellulitis and more life threating infections such as streptococcal toxic shock syndrome, necrotising fasciitis, and sepsis ^3,6^. The mechanisms that allow GAS to cause such diversity in disease types are unknown, however a number of studies have shown that bacterial and host-specific components may be involved ^7^.

GAS strains are typed based on the sequence of the *emm* gene, which encodes the M-protein, of which there are over 200 known *emm* types ^8^. The epidemiology of GAS infections has been changing globally over the last decade, with the emergence of new *emm* types and localised outbreaks a main feature ^9^. Within *emm* types of GAS, isolates may be causative of a range of clinical outcomes, such that most lineages carry the potential for expression of a range of phenotypes that may determine the course and nature of infection. Recent studies have shown distinct correlations between the host niche of recovered GAS clinical isolates and their ability to secrete high concentrations of known virulence factors such as streptococcal pyrogenic exotoxin A, B, and C (SpeA, SpeB, and SpeC), or the haemolytic exotoxin streptolysin O (SLO) ^10-12^. In addition to this, the premise that GAS phenotypic heterogeneity contributes to distinct clinical phenotypes is supported by studies that have found changes in virulence factor production such as in streptokinase and capsular protein secretion after GAS is passaged either *ex vivo* or *in vivo* ^13-17^.

The haemolysin SLO is well established as having cell and tissue destructive activity, and is part of the family of cholesterol dependent cytotoxins that also includes perfringolysin, pneumolysin, and listeriolysin O ^18-20^. SLO is a highly conserved protein secreted by nearly all clinical isolates of GAS, and acts against a wide number of eukaryotic cell types including macrophages, neutrophils, and erythrocytes, by interacting with cholesterol in target cell membranes to form pores, with sufficiently high doses of SLO resulting in complete cell lysis^5,21-23^. SLO has a number of other biological effects on the host that act at different stages throughout infection, such as its ability to cause hyper-stimulation and cell-meditated apoptosis of host immune cells such as neutrophils ^11,24^. Although most GAS isolates have the gene for SLO, the production of SLO is heavily regulated, as shown in studies that have seen variation in cytotoxicity within and between *emm* types ^25^. Early studies with SLO demonstrated that the purified toxin was lethal to mice and rabbits when injected intravenously, mainly due to cardiotoxicity ^26,27^. More recently, there have been studies to assess the effects of biologically relevant concentrations of SLO in *in vivo* models. Limbago *et al.*, found that SLO-deficient GAS resulted in attenuated skin infections and similarly Zhu *et al.*, reported a reduction in virulence when using SLO-deficient GAS in an invasive wound infection model ^28,29^.

Despite all this however, substantial gaps still exist in our understanding of the contributory role of the variation and amount of SLO production to overall GAS pathogenesis. In order to address this, we developed a custom made ELISA to compare the production of SLO between a recently emerged hypervirulent outbreak strain (which resulted in an epidemic in Liverpool, UK, between 2010-2012) characterised as *emm* type 32.2 and invasive *emm* type 1.0. isolates^30^. Using *in vivo* GAS bacteraemia and novel septic arthritis models, we further investigated the role of SLO in establishing and maintaining different clinical pathotypes *in vivo*. In addition, we investigated the role of SLO in these models, by use of a SLO deficient mutant strain in the background of the invasive outbreak *emm* type 32.2 isolate.

## Results

### *In vitro* phenotypic analysis of invasive and non-invasive *emm* type strains

Capsule thickness, complement deposition, and opsonophagocytic killing by macrophages was measured across 24 individual GAS isolates, covering 4 different *emm* types. Differences in capsule thickness, complement deposition, and opsonophagocytosis varied between and within *emm* type isolates. In general, invasive *emm* type 32.2 isolates had thicker capsules than non-invasive isolates (regardless of *emm* type) (Figure 1A) and had less complement deposited on their surface (Figure 1B). The size of the capsule and complement deposition were inversely proportional, with thinner capsule isolates exhibiting more complement deposition on their surface. The Pearson correlation coefficient (r) after log_10_ transformation was −0.5845 (r^2^ = 0.3416; p = 0.0027). The mean percentage of killing by macrophages in *emm* type 32.2 isolates (30% killing) was marginally lower than non *emm* type 32.2 isolates (37% killing) (Figure 1C). Although there was no significant correlation between capsule thickness and opsonophagocytosis, there was a positive association between the two (r = 0.20).

**Figure 1.**
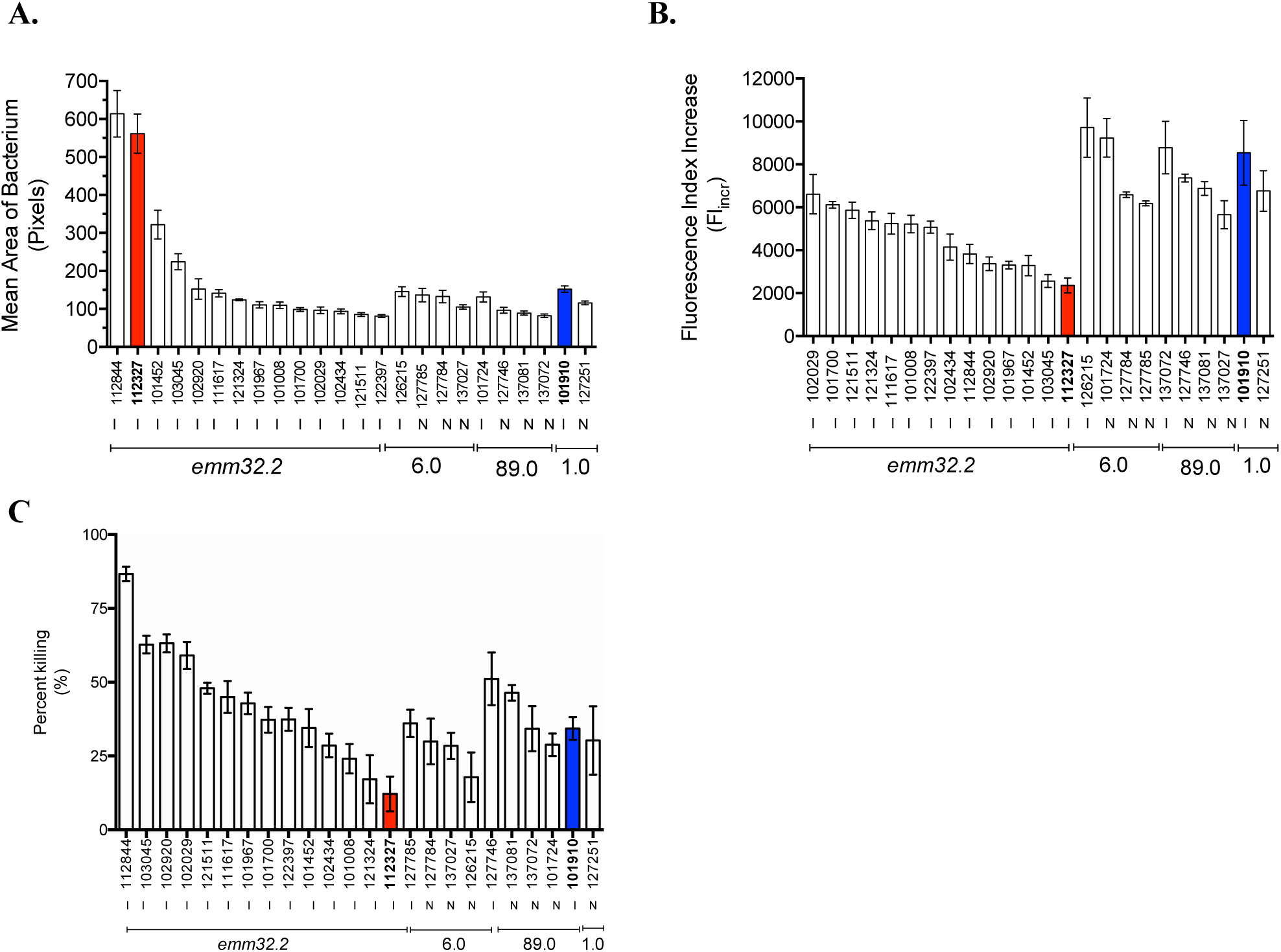
*In vitro* characterisation of capsule thickness, complement deposition, and opsonophagocytic killing of invasive and non-invasive *emm* type strains. A) Capsule thickness assay of 24 GAS isolates from 4 different *emm* types. B) Complement deposition assay. C) Opsonophagocytic killing assay. Isolates were analysed in triplicate and in three independent experiments for each assay, values are presented as mean ± S.E.M. I, invasive; N, non-invasive. Red and blue columns denote isolates selected for future comparison in mouse model experiments.

Based on these results, we chose representative isolates from the *emm* type 32.2 outbreak strains (isolate 112327) and *emm* type 1.0 strains (isolate 101910) which were most significantly different in capsule thickness, complement deposition and phagocytic killing scales, to use in subsequent *in vivo* infection modelling. As such, both isolates were from an invasive clinical phenotype but had significantly distinct phenotypic differences, with the e*mm* type 32.2 isolate 112327 exhibiting a significantly thicker capsule, lower complement deposition and more resistance to killing in comparison to the *emm* type 1.0 isolate 101910.

### *In vivo* characterisation of *emm* type 1.0 and 32.2 isolates in models of invasive GAS infection

An invasive GAS model was used to compare the virulence of *emm* type 1.0 (isolate 101910) and *emm* type 32.2 (isolate 112327) *in vivo.* Following intravenous infection, 100% of mice infected with isolate 112327 at 10^8^ CFU succumbed to infection by 24 h post-infection. When infected with a ten fold lower dose of 10^7^ CFU, all mice showed signs of lethargy by 24 h and succumbed to infection by 36 h post-infection (Figure 2A). In contrast, none of the mice infected with either 10^8^ or 10^7^ CFU of isolate 101910 showed any signs of infection and all survived (Figure 2A). In time point experiments over 48 h, mice infected with 10^8^ CFU of isolate 112327 had significantly higher bacterial loads in the blood at all time points post infection in comparison to those infected with isolate 101910, this was also the case by 24 h for isolate 112327 infected at tenfold lower (10^7^) CFU dose, suggesting that overall, *emm* type 32.2 (isolate 112327) is better adapted to survival and proliferation in blood.

**Figure 2.**
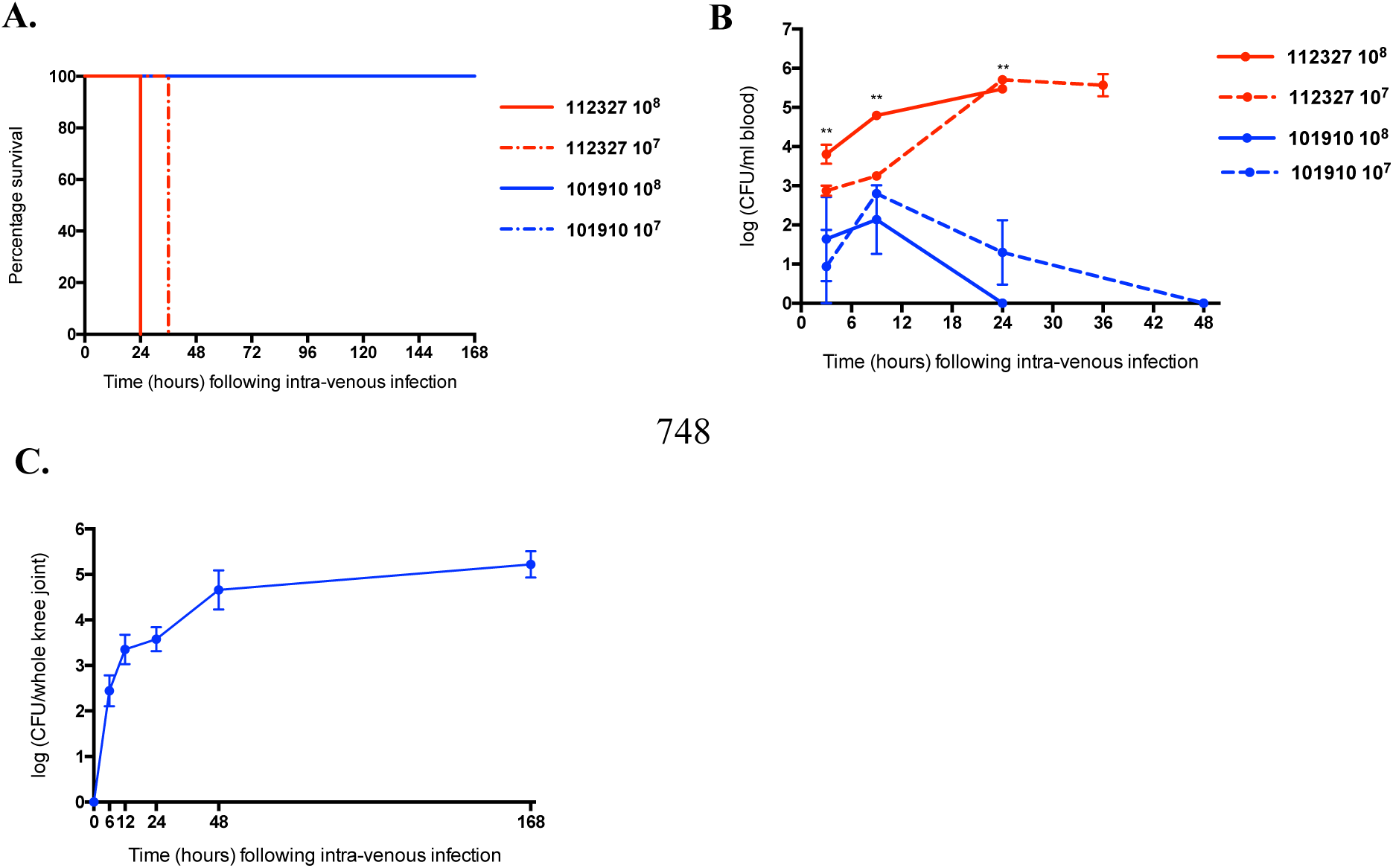
*In vivo* characterisation of *emm* type 32.2 (isolate 112327) and *emm* type 1.0 (isolate 101910) in a model of invasive GAS infection. A) Kaplan Meier plots representing percentage survival of CD1 mice (n = 10 per group) following 10^8^ and 10^7^ CFU intravenous infection with *emm* type 1.0 (isolate 101910) and *emm* type 32.2 (isolate 112327). B) The bacterial CFU in blood for each isolate and infectious dose over time. C) The bacterial CFU in knee joints of CD1 mice (n = 10, knee joints n = 20) following intravenous infection with 10^7^ CFU (50 µl) of *emm* type 1.0 (isolate 101910). **p-value<0.01 when analysed using a one-way ANOVA followed by a Kruskall-Wallis multiple comparisons test.

There was no detectable CFU of isolate 101910 in blood by 24 h (at 10^8^ dose) and by 48 h (at 10^7^ dose), demonstrating a clear difference in blood survival (Figure 2B) suggesting that *emm* type 1.0 (isolate 101910) was either less well adapted to survive in blood or was able to rapidly translocate elsewhere. Indeed, mice infected with isolate 101910 began to show symptoms of joint deformities by 24 h, which progressed until the end of the experiment. Bacteria were recovered from the knee joints at a mean log 2.4 CFU/knee joint as early as 6h (Figure 2C), and the bacterial load continued to increase up to a mean log 5.2 CFU/knee joint by the end of the experiment (day 7) (Figure 2C).

### Comparison of SLO production and activity in *emm* type 1.0 and 32.2 isolates

To explain the clear differences in overall mouse survival, bacterial virulence and proliferation in blood between the two *emm* type 1.0 and 32.2 isolates, we quantified the amount of SLO secreted by each isolate *in vitro* (at equivalent CFU) by analysing the amount of SLO directly secreted into the supernatant by the bacteria during growth phase in planktonic culture. The concentration (ng/ml) of SLO produced by *emm* type 32.2 (isolate 112327) increased rapidly over time compared with *emm* type 1.0 (isolate 101910); whereby isolate 112327 produced significantly more SLO from 6 h onwards until the final time point at 12 h (p = 0.015 - <0.0001). Isolate 101910 produced a small amount of SLO initially but the concentration did not continue to increase beyond 8 h (Figure 3A). We next looked at the haemolytic activity of SLO. The haemolytic activity of SLO secreted by isolates 112327 and 101910 followed the same pattern as that of the amount secreted; isolate 112327 SLO was significantly more haemolytic from 6 h until the final time point at 12 h compared to isolate 101910 SLO (p = 0.028 - <0.0001) (Figure 3B). Hence, *emm* type 32.2 (isolate 112327) secreted not only significantly more SLO than *emm* type 1.0 (isolate 101910), but also significantly more haemolytic toxin at equivalent CFU. The CFU growth of both isolates was assessed to ensure the differences observed for SLO were not due to significant differences in bacterial growth. We found no significant difference in bacterial growth for both isolates across all time points, with almost identical CFU loads at 10 and 12 h (Figure 2C), which interestingly were the same time points with the greatest difference in SLO concentration and activity, suggesting that bacterial growth rate and CFU load were not responsible for observed SLO differences between isolates.

**Figure 3.**
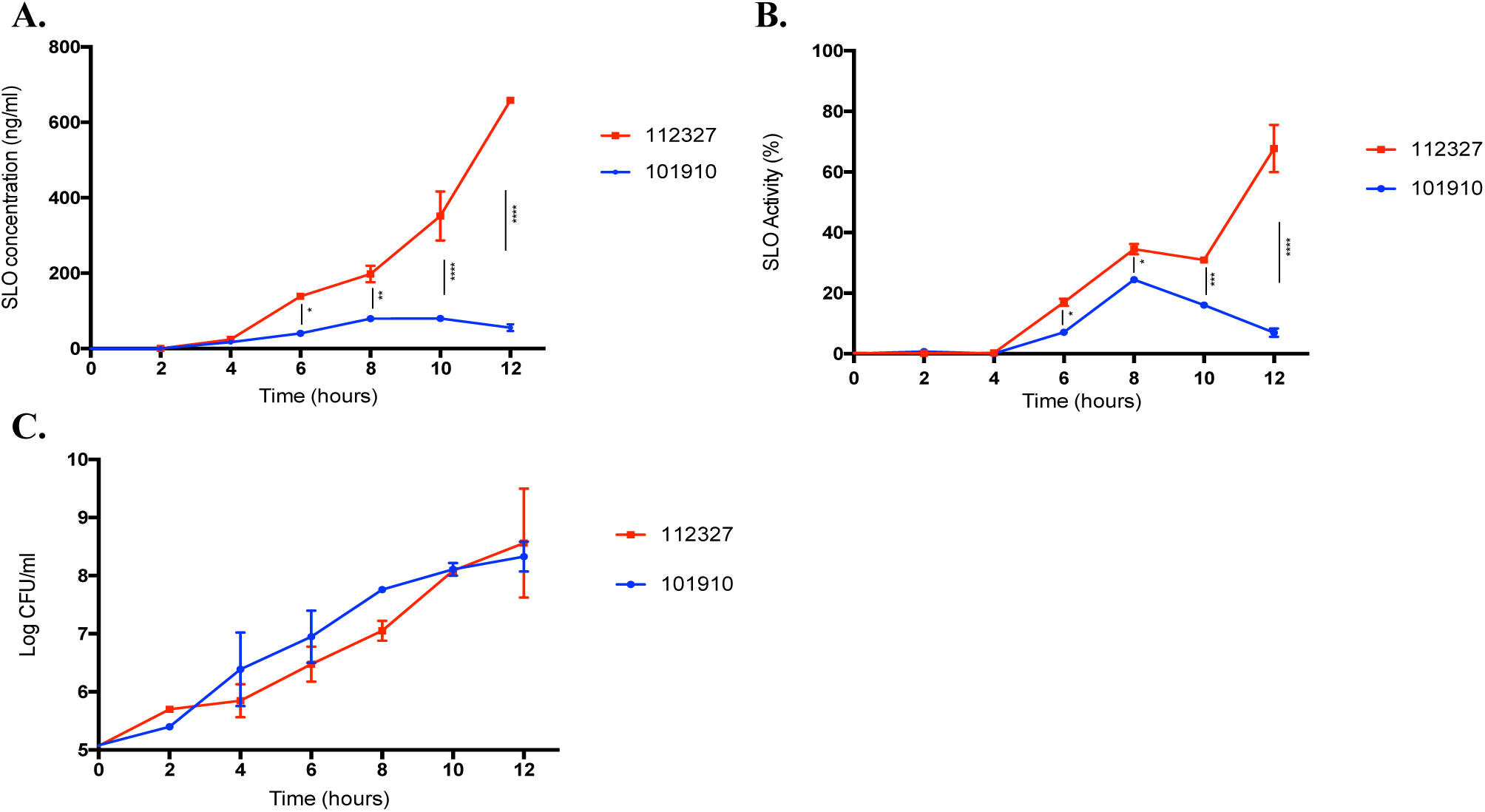
Comparison of streptolysin production and activity in *emm* type 32.2 (isolate 112327) and *emm* type 1.0 (isolate 101910). A) Concentration of streptolysin (ng/ml) secreted into the supernatant by *emm* type 32.2 (isolate 112327) and *emm* type 1.0 (isolate 101910) over time, measured by a custom made SLO-ELISA. B) SLO haemolytic activity C) and growth kinetics of isolates displayed as CFUs. *p-value<0.05, **p-value<0.01,***p-value<0.005, and ****p-value<0.001 when analysed using a two-way ANOVA followed by a Bonferroni’s multiple comparisons correction.

### *In vivo* recovered *emm* type 1.0 (isolate 101910) has reduced production and activity of SLO

To further clarify the reasons for *emm* type 1.0 (isolate 101910) clearance from bloodstream and translocation to the knee joints, we recovered bacteria from mouse knee joints at 24 h post infection, and quantified the amount of SLO secreted into the supernatant over an *in vitro* growth phase. We found that *in vivo* recovered isolate 101910 secreted significantly less SLO into the supernatant over the 12 h *in vitro* growth phase. There was significantly less SLO secreted from 6 h onward to that originally produced by isolate 101910 grown *in vitro* at equivalent CFU (p = 0.009 – <0.0001) (Figure 4A). In addition to producing significantly less SLO, the haemolytic activity of *in vivo* recovered bacterial SLO was also significantly lower from 6 h to 10 h, again at equivalent CFU (p = 0.0007 – <0.0001), with SLO activity comparable at 12 h (Figure 4B). We next wanted to determine whether *in vivo* recovered isolate 101910 retained its low SLO production and low SLO activity phenotype when grown over multiple times *in vitro*. Interestingly, after just one growth phase in THYG culture, the concentration of SLO reverted to a high SLO production phenotype (p = <0.0001) (Figure 4C), with significantly increased haemolytic activity from 6 h to 10 h (p = <0.005) (Figure 4D), suggesting that factors *in vivo* caused isolate 101910 to suppress its SLO production. During intravenous infection with *in vivo* recovered isolate 101910 (10^7^ CFU), bacteria enter the knee joint at higher bacterial numbers and proliferate in the joint more quickly. These results suggest that *in vivo* recovered *emm* type 1.0 (isolate 101910) is more adapted to initiating an infection in the joint more quickly.

**Figure 4.**
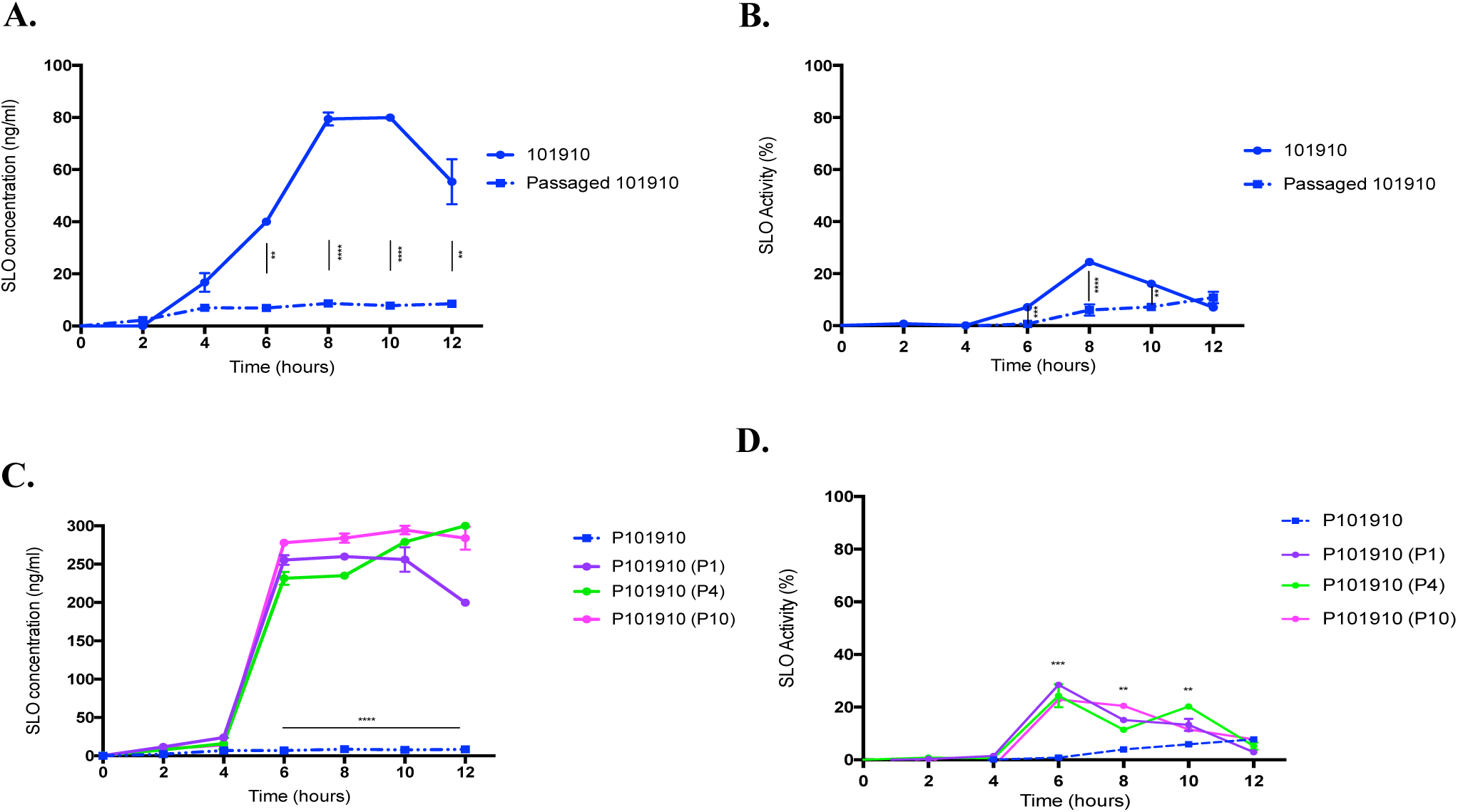
Comparison of streptolysin production and activity in *emm* type 1.0 (isolate 101910) grown *in vitro* or recovered from *in vivo*. A) Concentration of streptolysin (ng/ml) secreted into the supernatant by *emm* type 1.0 (isolate 101910) grown *in vitro* or recovered from knee joints (P101910) and then grown *in vitro*. B) SLO haemolytic activity. C) After subsequent *in vitro* passaging of *in vivo* recovered P101910 in Todd-Hewitt broth, the concentration of streptolysin (ng/ml) was measured, D) and the SLO haemolytic activity. **p-value < 0.01, ***p < 0.005 and ****p-value < 0.001 two-way ANOVA followed by a Bonferroni’s multiple comparisons correction.

### Concentration and activity of secreted SLO has significant impact on virulence *in vivo*

To investigate the effect of secreted SLO on virulence *in vivo*, we tested the amount and activity of SLO released into the challenge inoculum (prior to infection of mice) of *emm* type 1.0 (isolate 101910) and *emm* type 32.2 (isolate 112327). At equivalent CFU challenge inoculum (1 × 10^8^ per 50 μl), isolate 112327 had significantly higher SLO concentration (p = 0.012) (Figure 5A) and haemolytic activity (p = 0.01) (Figure 5A) than isolate 101910. This had a direct effect on survival *in vivo*, where mice infected with isolate 112327 all died from their infections, while those infected with isolate 101910 all survived (Figure 5C).

**Figure 5.**
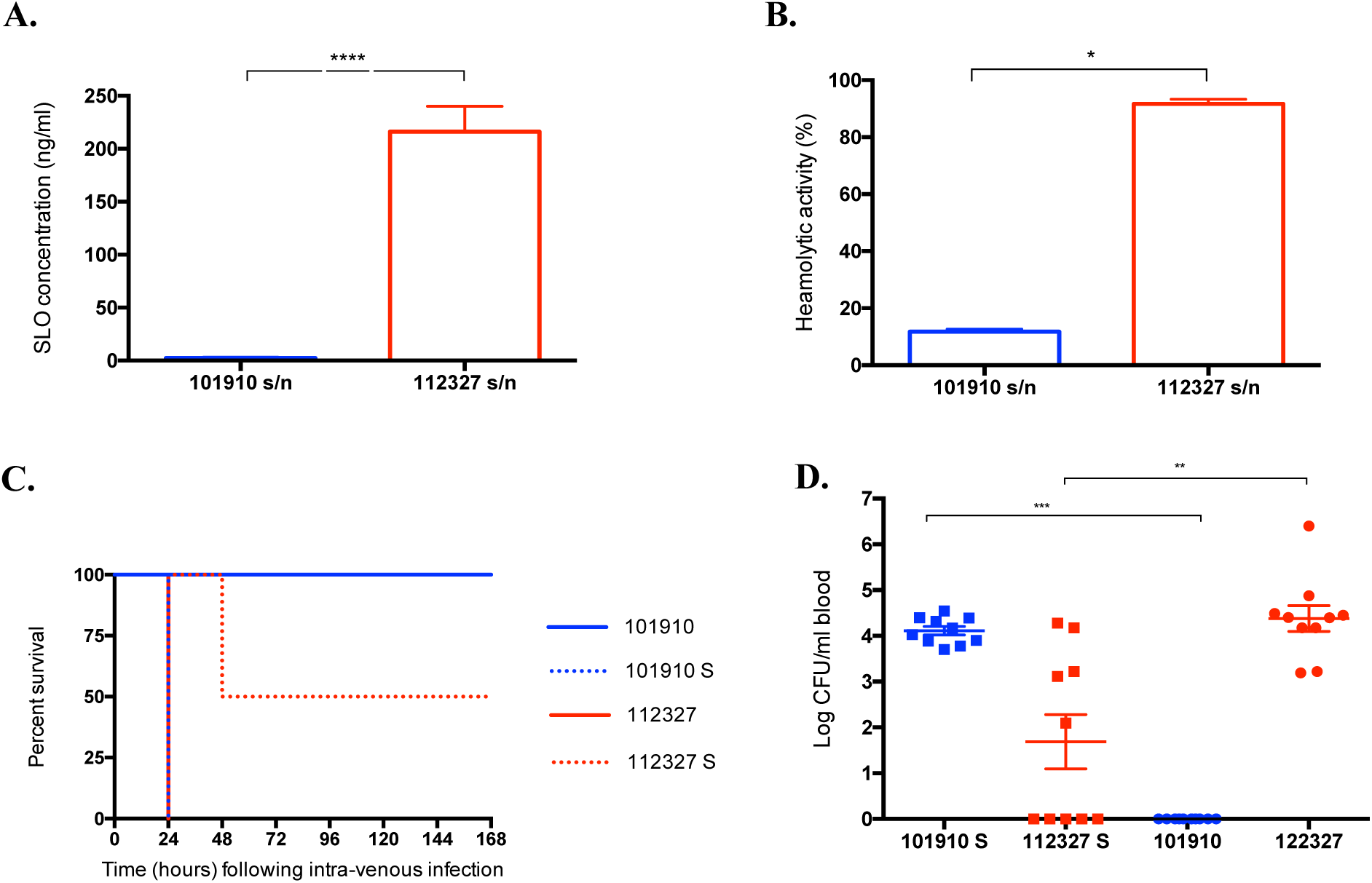
Effect of concentration and activity of secreted SLO on virulence *in vivo*. A) Concentration of streptolysin (ng/ml) and B) haemolytic activity in infection doses of *emm* type 1.0 (isolate 101910) and *emm* type 32.2 (isolate 112327), when prepared in 1 ml of PBS, incubated at room temperature for 30 minutes. C) Kaplan Meier survival plots representing survival of CD1 mice (n = 10 per group) when intravenously infected (10^8^ CFU) with isolates 101910 and 112327, and 112327 bacteria re-suspended in supernatant from 101910 challenge dose (112327 S) or 101910 bacteria re-suspended in supernatant from 112327 challenge dose (101910 S). D) Bacterial burden in blood 24 hours after infection with isolates 101910 and 112327 and swapped supernatant isolates as above. **p-value < 0.01, ***p < 0.005 and ****p-value < 0.001 when analysed using a one-way ANOVA followed by a Kruskall-Wallis multiple comparisons test.

To further investigate the effect of secreted SLO on infection dose and survival, the supernatants between isolates 112327 and 101910 were swapped prior to infection of the mice. Infection doses were prepared in 1 ml of PBS, incubated at room temperature for 30 minutes, immediately prior to infection bacteria were pelleted by centrifugation and the supernatants of the two challenge doses were swapped. Mice were infected with either isolate 112327 bacteria re-suspended in supernatant from isolate 101910 challenge dose or isolate 101910 bacteria re-suspended in supernatant from isolate 112327 challenge dose. In contrast to original challenge dose infections, the supernatant swap infected mice exhibited the opposite phenotype, whereby the normally non-lethal isolate 101910, now killed all mice when infected with supernatant from isolate 112327, and the normally lethal isolate 112327 isolate became less virulent, leading to only 50% death as compared to 100% death previously (Figure 5C).

Moreover, we determined the bacterial load in blood 24 h post-infection. As previously observed, there were no CFUs of isolate 101910 in blood at 24 h, but a significant 4 Log increase in CFUs was observed when isolate 101910 was infected with the supernatant swap dose, clearly suggesting that the high concentration of SLO present in isolate 112327 supernatant was enabling proliferation and retention of isolate 101910 in blood as compared to its normal condition of being cleared from blood (Figure 5D). In contrast, isolate 112327 challenge dose with isolate 101910 supernatant infected mice had significantly lower CFUs in blood at 24 h in comparison to when infected with its original supernatant (p = 0.0079) (Figure 5D) suggesting again, that the concentration and activity of SLO is key to virulence *in vivo*, both in terms of survival and bacterial load.

### SLO deficiency significantly reduces bacterial load and increases *in vivo* survival

To assess the role of SLO in the virulence of *emm* type 32.2 (isolate 112327) *in vivo* we generated an isogenic SLO deletion mutant where the SLO gene was deleted and replaced by a spectinomycin resistance gene through allelic exchange. All mice infected intravenously with 112327 ΔSLO mutant survived till the end of the experiment (96 h post infection), compared to mice infected with the wild type isolate whom all succumbed to infection by 24h (Figure 6A). The difference in mouse survival can be explained by the bacterial load in blood, which was 3.5 log lower in the ΔSLO mutant by 24h post infection, than that observed in mice infected with the wild type isolate (p = <0.0001) (Figure 6B). Furthermore, the bacterial burden of the 112327 ΔSLO mutant decreased over time and by 96 h post infection, bacteria were completely cleared from the blood (Figure 6B). These results indicate that in the absence of the toxin, bacteria were less able to establish an infection in the blood or were able to translocate elsewhere. Indeed, as described before during infection with *emm* type 1.0 (isolate 101910), mice infected with *emm* type 32.2 (isolate 112327) ΔSLO mutant also began to show joint deformities by 24 h. We detected a high bacterial load in the knee joints, contrary to mice infected with the wild type isolate in which no bacteria in knee joints or deformities were observed (Figure 6C). Interestingly, we detected no difference in the bacterial burden in the joints between 112327 ΔSLO mutant and *emm* type 1.0 (isolate 101910) (Figure 6C), which demonstrates that the kinetics of the infection was the same across the two different isolates and suggests that as well as being a key factor necessary and sufficient to virulence *in vivo*, SLO is also intriguingly the driving factor in determining the phenotypic outcome of infection.

**Figure 6.**
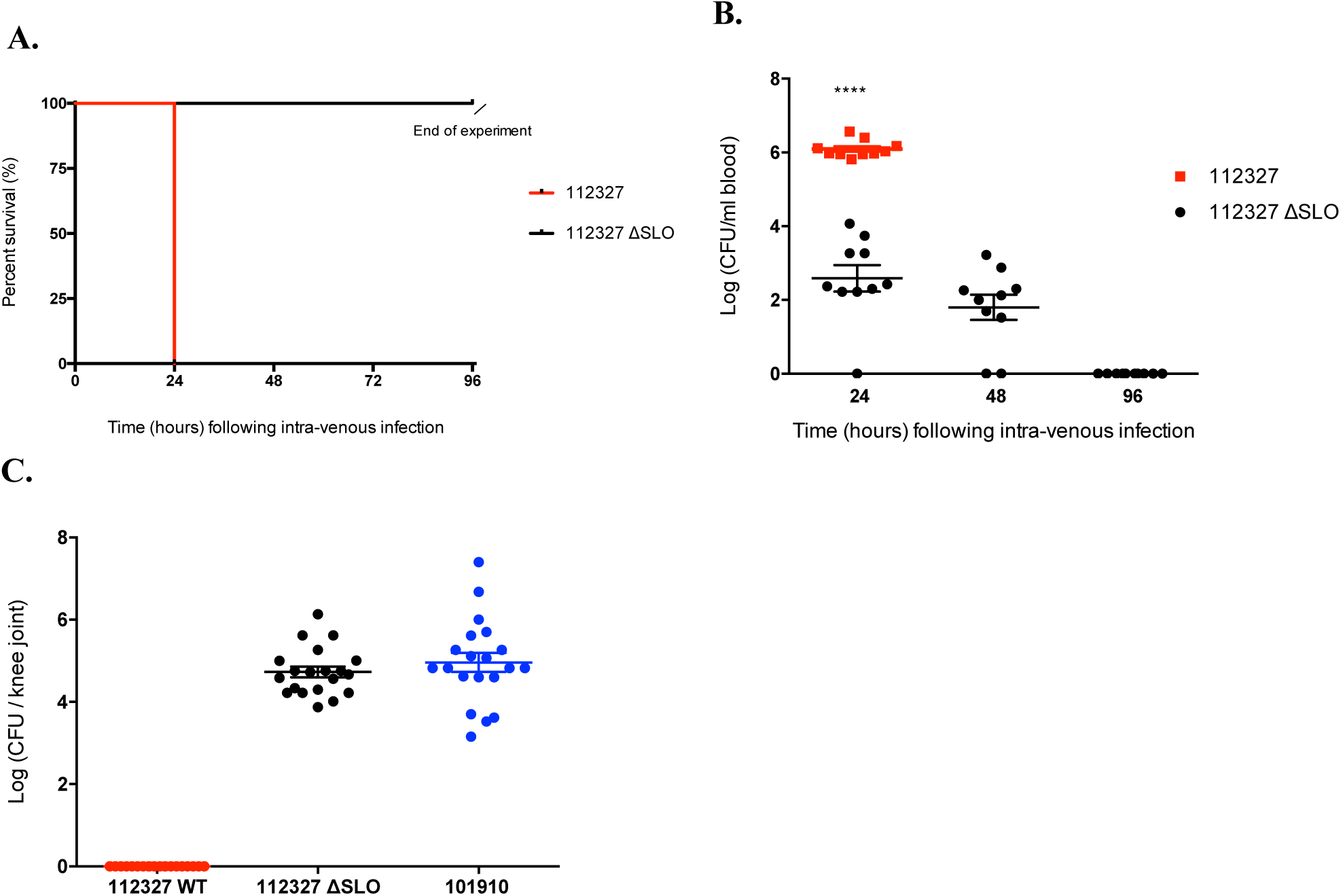
SLO deficiency increases *in vivo* survival and switches phenotype. A) Kaplan Meier plots representing percentage survival of CD1 mice (n = 10 per group) following 10^8^ CFU intravenous infection with isolates *emm* type 32.2 (isolate 112327) and *emm* type 32.2 (isolate ΔSLO 112327). B) The bacterial CFU in blood for each isolate over time. C) Bacterial load in knee joints (n = 20) at 24 h. ****p-value < 0.0001 when analysed using a two tailed Mann-Whitney U test.

### Liposome treatment reduces bacterial load and increases survival

Our previously published work has explored using cholesterol rich liposomes (cholesterol: sphingomyelin liposomes; 66 mol/% cholesterol) as a method to sequester cholesterol dependent cytolysins both *in vitro* and *in vivo* ^31,32^. We have shown that administration of cholesterol rich liposomes within 10 h after initiation of infection stopped the progression of bacteraemia caused by *S. aureus* and *S. pneumoniae* ^31^ and that liposomes were also able to bind strongly to SLO ^31,32^. Based on this, we now used specially tailored cholesterol rich liposomes as targets to sequester secreted SLO *in vivo*.

Liposomes were successful in sequestering the toxin as the concentration of SLO was significantly lower in *emm* type 32.2 (isolate 112327) supernatant when incubated with liposomes as compared to non-liposome control (p = 0.003) (Figure 7A). When the bacterial challenge dose was co-incubated with liposomes prior to infection, all mice infected with liposome treated isolate 112327, survived a further 24 h as compared to non-liposome treated control challenge dose (Figure 7B) and this extended survival period correlated with reduced CFU load in blood (Figure 7C). In addition, the effects of giving liposomes as a treatment to invasive GAS was also considered. A single injection of the liposomal mixture was administered at 4 h post infection. All mice that were not liposome treated succumbed to infection by 24 h, in comparison 60% of mice treated with the liposomal mixture survived an extra 24 h to 48 h (Figure 7D). In line with the differences in survival time, there was a considerable reduction of the bacterial load in the blood of mice that had been injected with a single dose of liposomes at 4 h in comparison to no treatment group (Figure 7E). These results show that a treatment with liposomes reduces the amount of SLO secreted into the extracellular environment, leading to a considerable reduction of the bacterial burden in the blood of mice and to attenuation of invasive GAS infection.

**Figure 7.**
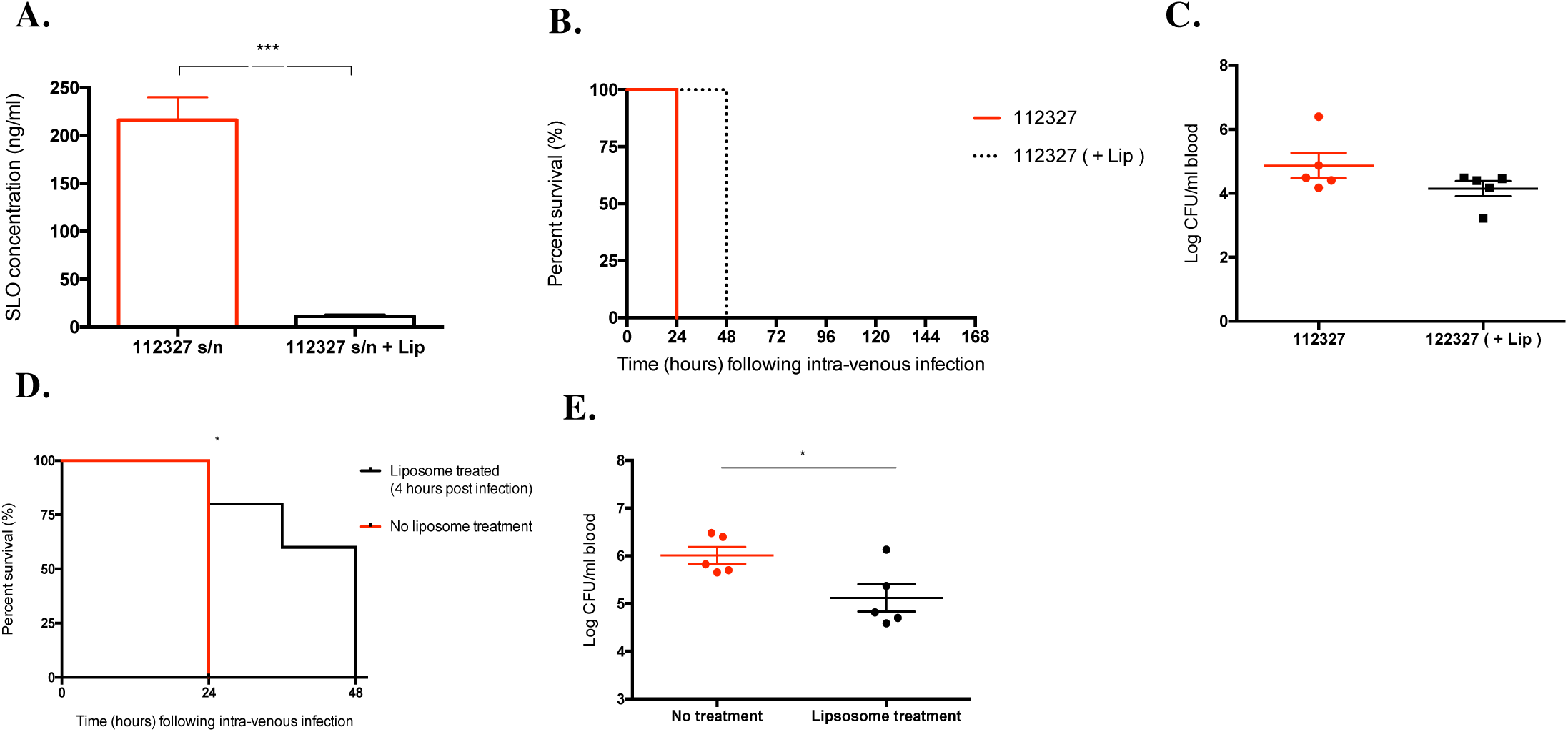
Liposome SLO treatment reduces bacterial burden and increases survival. A) Concentration of streptolysin (ng/ml) in infection doses of *emm* type 32.2 (isolate 112327) and *emm* type 32.2 (isolate 112327) after liposome treatment. B) Kaplan Meier survival plots representing survival of CD1 mice (n = 5) when intravenously infected (10^8^ CFU) with *emm* type 32.2 (isolate 112327) and after treatment of *emm* type 32.2 (isolate112327) supernatant with 4 µg/ml of liposomes. C) Bacterial burden in blood 24 hours after infection. D) Kaplan Meier survival plots comparison representing survival of CD1 mice (n = 5 per group) when intravenously infected (10^8^ CFU) *emm* type 32.2 isolate 112327 and after injection of liposomal mixture 4 h after infection. E) Bacterial burden in blood 24 hours after infection. Survival data was analysed using the Log-rank (Mantel-Cox) test (*p < 0.05). Data displayed as mean SEM and analysed using a two tailed Mann-Whitney U-test (*p < 0.05, ***p<0.005).

## Discussion

In this study, we investigated the role of SLO in determining disease phenotype, and found that differences in production and activity of SLO was central to *in vivo* pathotype and disease outcome in GAS infections.

In summary, we found that SLO production and activity drove two very distinct in vivo pathotypes; *emm* type 32.2 isolates which produced SLO in high levels and with high activity and *emm* type 1.0 isolates, which were the exact opposite with low levels of SLO production and of low activity. This correlated directly with their *in vivo* pathotype i.e. high virulence in bacteraemia models accompanied by short host survival (*emm* type 32.2) and low virulence in chronic septic arthritis models accompanied by long term host survival (*emm* type 1.0). Indeed, we found that the levels and activity of SLO at time of initial infection, determined the disease phenotype, with high levels of SLO driving invasive disease and low levels sustaining chronic joint infections. When removing SLO from the *in vivo* environment, either by gene deletion or by significantly reducing SLO (by supernatant swap or liposome sequestration method), we were able to demonstrate a complete reversal in the *in vivo* pathotypes of these *emm* isolates, whereby normally bacteraemia causing *emm* type 32.2 isolates could be made to translocate into joints rather than killing their hosts, and septic arthritis causing *emm* type 1.0 isolates could be made highly invasive, highlighting the crucial role of SLO in determining disease phenotype and outcome in vivo.

SLO is a major virulence factor for GAS, expressed by nearly all strains, and with amino acid sequence homology highly conserved between strains ^33^. Multiple roles in pathogenicity *in vivo* have been attributed to SLO, and recent studies have shown that SLO is important in the evasion of the host response via a number of mechanisms. Timmer *et al*., demonstrated that GAS induces rapid macrophage and neutrophil apoptosis due to the effects of SLO ^24^, and further work in the field has demonstrated that SLO rapidly impairs neutrophil oxidative burst preventing the bactericidal action of neutrophils ^34^. The effects of the general presence of secreted SLO in the blood stream has been less well studied however, although it has been implicated in driving inflammation, including the well documented evidence on activation of the NLRP3 inflammasome^35^, hence, it would therefore seem likely that SLO production and activity is important for GAS in invasive bacteraemia infections yet there has been no studies to date to show that SLO itself could be driving disease phenotype in vivo.

Although the SLO gene is highly conserved among all *emm* types of GAS, studies have shown that there are differences in the expression of the SLO gene which regulates the production of secreted SLO ^35^, and that specific invasive variants can be isolated post *in vivo* passage ^13^. In addition to this, it has been shown that *in vivo* conditions can result in differential expression of certain proteins; a study looking at exotoxins SpeA and SpeB found that *in vivo* host and/or environmental signals induced SpeA gene expression and suppressed SpeB expression that could not be induced under *in vitro* conditions ^15^.

This study demonstrates that SLO levels and activity determine invasiveness or chronicity during infection. The role of SLO was further investigated using supernatant switching, an SLO-deficient mutant and SLO sequestration by cholesterol rich liposomes. When the supernatant of isolate 112327 was replaced with isolate 101910 supernatant, the amount of SLO in the challenge inoculum was significantly reduced and 50% of the mice challenged were able to clear the infection, a delayed invasive phenotype was observed with mortality at 48 h instead of 24 h with 112327 and its original supernatant. This demonstrated that without the initial high SLO concentration in the challenge inoculum there is an attenuation of virulence.

The bacteria may secrete SLO during the infection but the initial challenge concentration remains the key determinant resulting in clearance when concentrations are low and increased virulence with higher concentrations. Interestingly, when we reversed this experiment and used the supernatant from the challenge inoculum of high SLO secreting isolate 112327 and co-infected that with isolate 101910, we saw a complete change in the clinical phenotype, whereby isolate 101910 was now able to successfully proliferate in the blood resulting in host death. Taking both of these results together, they indicate that the amount of SLO that is initially secreted is key to virulence in the early stages of infection, and it is possible for the host to successfully clear the bacteria when SLO concentrations are low. Our data also shows that the ability of the mice to survive infection is linked with the ability of the bacteria to proliferate. Early studies on SLO indicated that it was toxic when injected directly in an animal model ^28,36^, in our study administrating the supernatant alone into the mice without any bacteria did not have a fatal effect. The difference with our study could be due to early studies using supraphysiological concentrations of purified SLO and/or purified SLO preparations contaminated with LPS.

To consider how the complete removal of SLO affects the progression of invasive infection, an isogenic SLO deletion mutant of isolate 112327 was made. Our results demonstrate that mice infected with the SLO mutant had a significantly higher rate of survival than mice infected with its parent wild type bacteria. None of the mice infected with the mutant succumbed to infection where as 100% of mice infected with the wildtype died at 24 h. The SLO mutant began to be cleared from the blood as early as 24 h and was completely cleared by 96 h. Surprisingly, we found that the SLO deficient mutant sequestered in the knee joints causing septic arthritis as previously seen during infection with the low SLO secreting isolate 101910. The results clearly show that GAS strains lacking SLO and or low SLO producing GAS strains are severely impaired in their ability to cause bacteraemia and that lack (or reduced levels) of SLO enables the bacteria to accumulate within host joints.

There have been a number of previous studies using SLO mutants which have found that virulence is attenuated, although the relative importance of SLO would appear to be dependent on disease model used ^23,28,29,37^. For example, Limbago *et al*., used a subcutaneous invasive skin infection model to study the virulence of SLO-deficient mutants, where they found that although there were increased survival times of mice infected with SLO deficient strains, the absence of SLO itself did not limit dissemination from the wound into the vasculature ^28^. In contrast to this, a later study by Sierig *et al.*, found that during a skin infection model initiated by intraperitoneal infection there were no changes to survival using an SLO deficient mutant^23^. A more recent study looking at the emergence of an invasive *emm* type 89.0 clade, showed that elevated SLO producers are significantly more virulent than low SLO producers ^25^. Based on our findings here, we speculate that the low production of SLO (or SLO deficiency) prevents the ability of GAS to cause bacteraemia while enhancing its capability to translocate into the joints. Low SLO secreting isolate 101910 which effectively colonises the joints, adapts further to the joint by selecting for low secreting SLO variants. This is a selection pressure applied from environmental signals in the joint as when the isolate is recovered from the joints and placed under growth conditions *in vitro* it reverts to producing significantly more SLO (Figure 4C). Moreover, deletion of SLO in isolate 112327 resulted in a complete reversal of *in vivo* phenotype, whereby these SLO deficient isolates now caused septic arthritis.

The exact mechanism by how GAS infects the joint is not clear. The general mechanism of joint colonisation begins with haematogenous entry into the vascularised synovium. Once bacteria are in the joint space the low fluid shear conditions provide a unique opportunity for bacterial adherence and infection ^38^. Different strains of bacteria that commonly infect the joint including GAS and others such as *S. aureus* have varying degrees of tropism to the joint, thought to be due to differences in adherence characteristics and toxin production ^38^. We have previously shown that isolate 112327 is an outbreak strain with characteristics that suggest it is hypervirulent, e.g. it has 19 extra genes, five of which are associated with an increase in virulence ^39^. We have shown in this current study that isolate 112327 is more virulent in an invasive bacteraemia model and produced significantly more SLO which is likely to be one of the causes of its increased capacity to cause host death.

Infection with isolate 112327 results in uncontrolled bacterial proliferation in blood and rapid progression into sepsis. On the other hand, isolate 101910 which was isolated from a patient with septic arthritis and produces low concentrations of SLO, can be reduced to even lower concentrations after further selection from the joint. This implies that decreased production of SLO is beneficial during infection in the joint. Reduced or no expression of SLO could have a protective effect for the pathogen, as SLO is immunogenic and avoiding host immune cell detection could thereby prevent immune activation and clearance, allowing GAS to continue to colonise the joint^21^.

The results presented here have important implications for our understanding of GAS pathogenesis. We conclude that levels and activity of SLO is key to determining whether GAS infection follows an highly invasive and virulent pattern leading to host death or whether it follows a chronic pattern of long term joint infection. The fact that these disease phenotypes are not fixed is highly interesting, as it suggest that GAS is highly sensitive to environmental signals and can change its phenotype rapidly. Indeed, by artificially effecting SLO levels, we have shown that one disease phenotype can easily be switched into another. This has significant implications for therapy and vaccines as anti-SLO based treatments may not be the complete answer to protection against all forms of GAS infection.

## Materials and methods

### Epidemiological study design and collection of isolates

As previously published in Cornick *et al.*, between January 2010 to September 2012, the Respiratory and Vaccine Preventable Bacteria Reference Unit (RVPBRU) in the United Kingdom confirmed a total of 14 cases of *emm* type 32.2 invasive GAS in the Merseyside area. Over the same time period, 30 non-*emm* type 32.2 invasive GAS infections were collected alongside 20 non-invasive pharyngitis GAS isolates supplied by the Royal Liverpool University Hospitals Trust and Alder Hey Children’s Hospital ^30^. The Merseyside outbreak *emm* type 32.2 isolates (n = 14) were selected for this study alongside a selection of non-*emm* type 32.2 isolates. Invasive (n = 2) and non-invasive (n = 2) of each *emm* type 6.0 and *emm* type 89.0 isolates were selected and an additional well studied invasive *emm* type 1.0 isolate. All isolates were stored in Microbank^™^ beads prior to study.

### Bacterial culture conditions

Isolates were routinely grown on blood agar base (Oxoid) supplemented with 5% fresh horse blood and incubated overnight at 37°C in a candle jar. Liquid cultures were prepared in Todd Hewitt broth with 0.5% yeast extract and 0.5% glucose (THYG) and grown overnight at 37°C. Stocks of GAS in exponential growth phase were prepared by inoculating THYG broth with overnight cultures (1:40), and incubating at 37°C for 3-4 h. Glycerol was added (20% v/v) and stocks were stored at −80°C.

### Measurement of capsular thickness

Capsule thickness was measured using the FITC-dextran zone of exclusion method, as previously described, with minor modifications ^40^. Exponential phase cultures were centrifuged at 3000 g for 10 minutes, and the pellet re-suspended in PBS. 10 µl of bacterial suspension was mixed with 1 µl of 2000 kDa FITC-dextran (Sigma-Aldrich) and pipetted onto a microscope slide. The Nikon Eclipse 80i fluorescence microscope (100x magnification) was used to view the slides and photographs were taken using a Hamamatsu C4742-95 camera. ImageJ was used to determine the zone of exclusion (area in pixels), a value proportional to capsular thickness.

### Complement deposition

The complement deposition assay was based on a previously published method ^41^. Briefly, bacteria was added to BHI broth, incubated at 37°C for 15 minutes, and centrifuged. The supernatant was removed and the pellet was washed and re-suspended for incubation in PBS with 20% human serum (pooled from five individuals) and 1% gelatin veronal buffer. After washing, they were re-suspended in mouse-anti-human-C3 in PBS (Abcam) and incubated at 37°C for 30 minutes. Washing was repeated, and the contents were re-suspended in anti-Mouse IgG2a-APC in PBS (EBioscience) and incubated at 4°C for 30 minutes in the absence of light. After washing, the remaining bacteria were re-suspended in PBS and incubated with thiazole orange (BD Cell Viability kit). Samples were acquired using the Accuri C6 flow cytometer (BD).

### Opsonophagocytosis killing assay

The ability of isolates to resist killing by macrophages was measured using an adapted protocol of a previously published opsonophagocytosis killing assay (OPKA) ^42^. J774.2 macrophage cell line was maintained, as per standard protocols ^43^. Bacteria (1×10^5^ CFU/ml) were opsonised with IVIg (1:4) in HBSS (plus Ca2+/Mg2+, 5% FBS) for 20 minutes at 37°C with shaking at 180 rpm. Next, 1×10^5^ J774.2 cells were incubated with 5×10^2^ CFU of opsonised bacteria and 10 µl of baby rabbit serum complement (37°C, 45 minutes, 180 rpm). The CFU count in each well was then determined. Percentage killing was calculated from CFU remaining compared to control samples without J774.2 cells.

### *In vivo* models of invasive GAS infection

Seven-week-old CD1 mice (Charles River) were intravenously injected with PBS containing either 10^7^ or 10^8^ CFU of GAS in exponential growth phase. Following infection, mice were monitored for physical signs of disease using a standard scoring system ^44^. CFU counts were performed on blood collected at time points by tail bleeding. Mice were humanely culled when they were scored ‘++lethargic’ and blood tissue was collected for CFU enumeration. To make passaged stocks two CD1 mice were infected IV with 10^7^ bacteria. The mice were monitored to ensure that 24 h following infection they were at least a score of 1 on the arthritic index, a scoring system, which evaluates the intensity of arthritis, based on macroscopic inspection. The mice were humanely culled, the knee joints removed and bacteria recovered to make bacterial stocks. *In vivo* experimental procedures were reviewed by the University of Liverpool Ethical and Animal Welfare Committee and carried out under the authority of the UK Home Office Animals Scientific Procedures Act 1986 (UK Home Office Project Licence number P86DE83DA).

### Streptolysin ELISA design and method

Prior to analysis samples were thawed at room temperature. Plates (R&D systems) were coated with 1 µg/well monoclonal SLO antibody (Abcam) in PBS (Peprotech) at 4°C overnight. Plates were washed at each step with Peprotech washing buffer. After blocking (Peprotech), samples were added to the wells and incubated for 2 h at room temperature. The plate was washed (×5) and incubated with rabbit IgG polyclonal anti-SLO antibody (Abcam) for 2 h. Anti-rabbit IgG alkaline phosphatase conjugate secondary antibody (Abcam) was diluted to 1:5000 in blocking buffer, and after washing, was added and incubated for 30 mins. After washing, alkaline phosphatase yellow liquid substrate (PNPP) was added and incubated for 30 mins in the dark, to stop the reaction 1 M Sodium Hydroxide (NaOH) was used. The plate was loaded on to a Multiskan Spectrum (Thermo) and the absorbance measured at 405nm. All ELISAs were carried out with control wells which had all reagents added except samples or diluted SLO. Duplicate samples of each time point was measured on a single plate and repeated independently. Each plate contained six two-fold dilutions of a known concentration of SLO. The results were analysed using Sigma Plot and a standard curve developed to generate concentrations in ng/ml.

### Haemolytic activity assay

The haemolytic activity of SLO in culture supernatant was measured as previously described, with minor modifications ^45^. Bacteria-free supernatants were incubated at room temperature for 10 minutes with 20 mmol/l of dithiothreitol (Sigma-Aldrich). Supernatant was aliquoted into two tubes; 25 µg of water-soluble cholesterol (inhibitor for SLO activity) was added to one. Both tubes were incubated at 37°C for 30 minutes, followed by the addition of 2% sheep erythrocytes/PBS suspension to each sample and further incubation at 37°C for 30 minutes. PBS was added to each tube the samples were centrifuged at 3000 × g for 5 minutes. Each sample was transferred to a 96-well plate and the OD_541nm_ was measured.

### Generation of *slo* detion GAS mutant

An isogenic SLO knockout mutant of strain *emm* type 32.2 isolate 112327 was constructed through double-crossover allelic replacement of SLO with aad9 (encoding spectinomycin resistance). Regions directly upstream and downstream of SLO (∼ 1000 bp each) were amplified by PCR using primers SLO112327-up-F and SLO112327-up-R, SLO112327-down-F and SLO112327-donw-R respectively, which introduced BamHI restriction sites into the PCR products. These fragments were stitched together in a second round of PCR using primers SLO112327-up-F and SLO112327-donw-R, generating a 2 kb fragment with a central BamHI site, which was then ligated into pGEM-T vector (Promega), generating pGEM-T-Δslo-2kb. The plasmid was transformed into *E. coli* DH5α competent cells (ThermoFischer Scientific). The aad9 gene was amplified by PCR using primers aad9-F and aad9-R. The PCR product was subcloned into pGEM-T-Δslo-2kb at the BamHI restriction site, generating pGEM-T-Δslo::aad9, which interrupted the slo fragment, providing a means of positive selection of transformants. The generated plasmid was transformed into *emm* type 32.2 isolate 112327 by electroporation as previously described^46^. Transformants were recovered on THY agar supplemented with spectinomycin (100 µg/mL) at 37°C in a candal jar for up to 72 h. SLO deletion was identified by PCR and the PCR products were sequenced to confirm authenticity of the insertion.

### *In vivo* invasive model-switching supernatant of isolates

Frozen bacterial stocks were thawed at room temperature and 10^7^ bacteria were prepared in 1 ml PBS. After 30 mins both strains were centrifuged at 14,000 × g for 2 mins, the supernatant from *emm* type 32.2 (isolate 112327) was used to re-suspend *emm* type 1.0 (isolate 101910) bacteria and the supernatant from *emm* type 1.0 (isolate 101910) was used to re-suspend *emm* type 32.2 (isolate 112327) bacteria. Mice were immediately infected. The supernatants were analysed using the SLO-ELISA to measure the amount SLO present in ng/ml. Mice were humanely culled when they were scored ‘++lethargic’ and blood tissue was collected for CFU enumeration.

### Liposomes

Liposomes were generated with cholesterol and sphingomyelin from egg yolk from Sigma and dissolved in chloroform at 100 and 50mg/ml respectively. Lipids were mixed together with cholesterol at 66 mol/% proportion and then evaporated with nitrogen gas for 30min. For Cholesterol: Sphingomyelin (Ch:Sm) large and small liposomes, the hydration was made by addition of PBS (ThermoFisher scientific) and incubated at 55°C for 30 mins with vortexing. To obtain small unilamellar particles, the liposome preparation was then subsequently sonicated for 30min at 4°C. To eliminate carboxyfluorescein, the preparation was diluted in PBS and applied to a Sephadex G-25 column in PD-10 (GE Healthcare). Particle concentration and size distribution of the liposomes generated were evaluated using the NanoSight NS300 instrument (Malvern, UK) and using Nanoparticle Tracking Analysis (NTA) software.

## Data analysis

Statistical analysis was carried out using the GraphPad Prism^®^ version 5 statistical package (GraphPad Software, Inc. http://www.graphpad.com). The statistical significance according to the p-values were summarised as follows: *p-value<0.05, **p-value<0.01,***p-value<0.005 and ****p-value<0.001.

## Author Contributions

NF and AK conceived, designed and supervised the study and contributed equally throughout. JC, MB, MA, SP, MP and GP performed experiments. WAP and GP provided reagents. JC, NF and AK analysed data. JC and AK wrote the paper with input from all authors.

## Competing interests

All authors: No potential conflicts of interest.

## The Paper Explained

The shifting epidemiology of Streptococcus pyogenes (GAS) infections globally over the past decade has been punctuated by *emm* type emergence and localised outbreaks of severe invasive disease. Within *emm* types there is diversity of possible clinical outcomes, however, the basis of these varied clinical phenotypes is not well understood. To address this question, we investigated the role of GAS virulence and its host interactions. We discovered that streptolysin (SLO) can control disease outcome towards either acute pro-inflammatory or chronic disease phenotypes. The specific importance of these results are significant as while neutralisation of SLO activity will reduce severe invasive disease, this carries a risk of the promotion of chronic inflammatory conditions such as septic arthritis.

